# Correlate: A Web Application for Analyzing Gene Sets and Exploring Gene Dependencies Using CRISPR Screen Data

**DOI:** 10.64898/2026.04.02.716070

**Authors:** Soham Deolankar, Fredrik Wermeling

## Abstract

CRISPR screen data provides a valuable resource for understanding gene function and identifying potential drug targets. Here, we present Correlate, a freely accessible web application (https://correlate.cmm.se) that enables exploration of the Cancer Dependency Map (DepMap) CRISPR screen gene effects, hotspot mutations, and translocation/fusion data across more than 1,000 human cancer cell lines. The application supports two main use cases: (i) analysis of user-defined gene sets (e.g. CRISPR screen hits) to identify functionally linked genes based on correlations while providing an overview based on essentiality or user-provided screen statistics; and (ii) exploration of genes of interest in defined biological contexts, such as specific cancer types or mutational backgrounds, to generate hypotheses about gene function and dependencies. Additionally, Correlate supports experimental design by providing rapid overviews of gene essentiality and enabling the identification of cell lines with relevant mutational profiles. In contrast to knowledge-based approaches such as STRING and GSEA, which rely on prior biological annotations and curated interaction networks, Correlate identifies gene connections directly from functional CRISPR screen readouts, offering a complementary and data-driven perspective on gene network analysis. The application runs entirely in the browser, requires no installation or login, and integrates with the Green Listed v2.0 tool family for custom CRISPR screen design.

**HIGHLIGHTS:**

▪ Interactive web-based platform for bulk correlation analysis of user-defined gene sets using DepMap CRISPR screen data, requiring no installation or programming expertise.
▪ Identifies functional gene relationships from CRISPR screen readouts rather than curated annotations, offering a data-driven complement to tools such as GSEA and STRING.
▪ Enables contextual exploration of gene dependencies across cancer types and mutational backgrounds, supporting hypothesis generation about gene function and therapeutic targets.
▪ Supports experimental design through gene essentiality overviews, mutation and fusion analysis, and cell line identification, with optional integration of user-provided statistics from CRISPR screens, proteomics, or transcriptomics analyses.

## INTRODUCTION

Genome-wide CRISPR screens have become a central approach for systematically identifying gene dependencies in cancer (1,2,3). The Cancer Dependency Map (DepMap) project has performed CRISPR knockout viability screens across more than 1,000 cancer cell lines, generating gene effect scores that quantify how the inactivation of each gene affects cell fitness (4,5). The DepMap portal (https://depmap.org) additionally integrates mutational, gene expression, and drug response data, and has become an important resource for cancer biology, supporting the identification of potential therapeutic targets, the discovery of functional gene relationships, and the characterization of genotype-specific vulnerabilities (6).

An important insight from these large-scale datasets is that genes with correlated gene effect profiles across cell lines often share functional relationships (7-11). Identifying such correlations can reveal functional connections between genes, suggest pathway membership, and guide experimental design. Importantly, because these correlations are derived from functional readouts (based on cell survival and proliferation) rather than curated databases, they can complement more knowledge-based approaches such as gene set enrichment analysis (GSEA) and protein interaction databases like STRING (12,13). The relationships identified through this type of analysis are therefore particularly useful for functional interpretation and hypothesis generation.

The DepMap portal provides a range of tools for exploring these data, including co-dependency analysis for individual genes (4). Several additional tools have been developed to leverage DepMap data, including shinyDepMap for browsing gene essentiality and functional connections (14) and DepLink for linking genetic and pharmacologic dependencies (15). However, exploring functional relationships within user-defined gene sets, for example from a CRISPR screen, proteomics, or transcriptomics study, currently requires programming, as no dedicated web application supports this directly.

To address these limitations, we developed Correlate, a web-based tool for functional exploration of gene dependencies using DepMap CRISPR screen data. Correlate supports two primary analytical workflows: gene set-centered analysis to identify functional relationships and prioritize candidate genes, and gene-centered exploration to generate hypotheses about context-specific dependencies across cancer types and genetic backgrounds. By enabling bulk analysis and interactive network exploration, Correlate complements the DepMap portal and related tools. The tool is part of the Green Listed tool family (16,17) and extends previously described DepMap correlation workflows (17). Correlate is freely available at https://correlate.cmm.se.

## METHODS

### Development

Correlate was developed using JavaScript, HTML, and CSS without external frameworks or libraries. The application runs entirely in the browser with all computations performed client-side. Network visualization is rendered using the HTML5 Canvas API with a force-directed layout algorithm. Scatter plots and gene effect charts are generated dynamically. The application is hosted on an Apache HTTP server. Pearson correlation coefficients and Welch’s t-test statistics are computed using custom JavaScript functions. The source code is available at https://github.com/fredrikwermeling/correlation-web-app-.

### Data sources

Correlate uses a subset of data from the DepMap public release (currently 25Q3) (4), specifically the CRISPRGeneEffect.csv file containing gene effect scores processed using the Chronos algorithm (18), the Model.csv file with cell line annotations (lineage, subtype), the OmicsSomaticMutationsMatrixHotspot.csv file for hotspot mutation data, and the OmicsFusionFiltered.csv file for translocation/fusion data. Gene effect scores represent the fitness effect of gene inactivation, where a score of -1 corresponds to the median effect of known common essential genes and 0 corresponds to no effect.

### Correlation analysis

Pearson correlation coefficients are computed between gene effect profiles across available cell lines. Users can set a correlation cutoff (default 0.5), a minimum slope threshold, and the minimum number of cell lines required for a correlation to be reported. Correlations are visualized as an interactive network where edge thickness scales with correlation strength and edge color indicates positive or negative correlation.

### Mutation and fusion analysis

Mutation Analysis mode identifies differential genetic dependencies by comparing gene effect scores between cell lines with and without specific hotspot mutations in a target gene. P-values are calculated using Welch’s t-test, and effect sizes are reported as the difference in mean gene effect. Users can stratify by lineage and mutation count (0, 1, or 2 hotspot mutations). An equivalent analysis mode is available for translocation/fusion events.

## RESULTS

### Overview of Correlate

Correlate provides an interactive web-based interface for exploring gene effect correlations and dependencies in the DepMap CRISPR dataset. The application is organized around two main input areas: a parameter panel (“1. Set Parameters”), where the analysis mode, correlation cutoff, and filters are defined, and an input panel (“2. Input Genes”), where gene symbols are entered (**Fig. 1A**). Several analysis modes are available, including Analysis (correlations within the input gene set), Design (correlations between input genes and all other genes), Mutation Analysis (identifying differential gene effect by mutation status), and Synonym/Ortholog Lookup. After the user initiates the analysis, results are displayed as an interactive network graph with accompanying data tables and stratification options based on cancer type or hotspot mutations. Together, these modes provide a flexible framework for both exploratory analysis of user-defined gene sets and hypothesis generation in biologically defined contexts.

**Figure 1.**
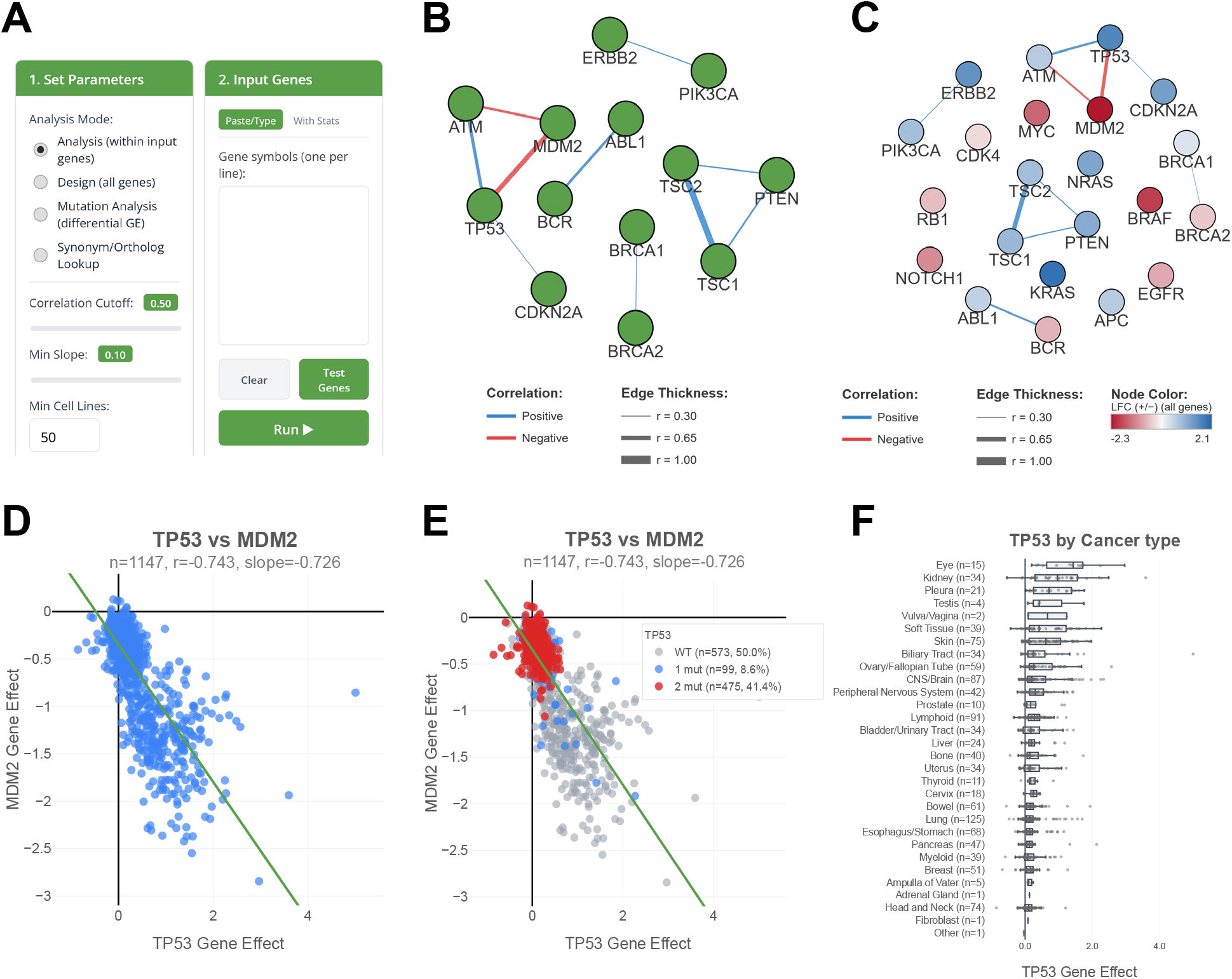
Gene set analysis in Correlate. (**A**) Screenshot of two key user interface elements: “1. Set Parameters” and “2. Input Genes”. (**B**) Correlation network of the default test gene set at a correlation cutoff of 0.3, showing functionally linked genes. (**C**) Network from the “With Stats” test gene set at correlation cutoff 0.3, with user-provided CRISPR screen statistics (log fold change) overlaid as node colors, allowing a rapid overview of screen results. (**D**) *TP53* vs *MDM2* scatter plot, accessed by double-clicking the connecting edge in the network. (**E**) *TP53* vs *MDM2* scatter plot with additional color stratification based on *TP53* hotspot mutation status. (**F**) *TP53* gene effect stratified by cancer type, accessed by double-clicking the node in the network. B-F are exported directly from the app as SVG files.

### Gene set analysis

In the default Analysis mode, Correlate identifies pairwise correlations within a user-provided gene set, using all available cell lines or a filtered subset. This allows exploration of functional relationships among user-provided hits from e.g. a CRISPR screen or functional redundancies among genes in a pathway of interest. The analyses can furthermore be filtered to a specific cancer type and/or cells with specific combinations of mutations. Significant correlations are displayed as a network (**Fig. 1B**), where edges connect genes whose inactivation has related fitness effects across cell lines. This provides a rapid overview of whether a candidate gene set contains coherent functional modules.

The correlation cutoff can be adjusted by the user; a cutoff of 0.3 to 0.5 is generally suitable for exploratory analysis, whereas higher cutoffs highlight stronger relationships. Correlate also reports and allows applying a cutoff to the slope of each pairwise relationship, as a strong correlation with a very low slope may reflect a statistically robust but biologically less meaningful association. An example of a biologically meaningful relationship recovered by this type of analysis is the strong inverse correlation between *TP53* and *MDM2* (r=-0.74), consistent with the well-established role of MDM2 as a negative regulator of TP53 function (10,11,19). Similarly, *TSC1* and *TSC2*, which encode the two components of the tuberous sclerosis complex (20), show one of the strongest positive correlations (r=0.91) in the dataset, consistent with their obligate functional interaction as a heterodimeric complex (21).

When gene lists are entered using the “With Stats” mode (under “2. Input Genes” in Fig. 1A), users can include quantitative values from gene-level analyses, such as log fold change (LFC) and FDR from CRISPR screens, for example as output by MAGeCK (22), or differential gene expression analyses. These values can be overlaid as node colors on the correlation network using the “Color by Stats” option (**Fig. 1C**), providing an integrated view of both the user-provided statistics and the functional relationships among hits. Alternatively, selecting “Color by GE” overlays DepMap gene effect scores as node colors, providing an overview of gene essentiality across the hit list. When combined with the option to display all input genes in the network, both modes provide a rapid overview of the full hit list that can inform downstream decisions, for example during the design of a custom CRISPR screen library. A summary table that includes correlation cluster assignments, DepMap gene effect scores, and user-provided statistics is available for download under the “Clusters” tab, providing a more manageable format for browsing large gene sets where the network visualization may become dense.

Double-clicking an edge in the network generates a scatter plot showing the pairwise gene effect scores across all the filtered cell lines. **Figure 1D** shows the *TP53* vs *MDM2* scatter plot: each data point represents a cell line, and the negative linear relationship illustrates that cell lines sensitive to *TP53* inactivation tend to be insensitive to *MDM2* inactivation, and vice versa, consistent with the established role of MDM2 as a negative regulator of TP53. This is the basis of the correlation analysis: genes with related gene effect profiles across cell lines may share functional relationships, whether through positive co-dependency or inverse association. A third dimension can also be added to the scatter plot by overlaying hotspot mutation status, enabling exploration of how mutational context influences gene dependencies. This is exemplified in **Figure 1E**, which shows that cell lines harboring TP53 hotspot mutations exhibit reduced sensitivity to both *TP53* and *MDM2* inactivation. Individual cell lines can also be identified and labeled in the scatter plot by clicking data points or by searching for specific cell lines in the application, supporting cell line selection for experimental follow-up. The settings bar allows filtering by cancer type, subtype, and mutation status. The “Compare by” buttons can further help identify which filters yield the strongest correlations. Double-clicking a node in the network displays the gene effect of the selected gene stratified by cancer type (**Fig. 1F**), providing a rapid overview of which lineages are most dependent on the gene.

### Hypothesis generation

The Design mode (selected in “1. Set Parameters”, indicated in Fig. 1A) enables hypothesis generation by identifying genome-wide correlates of one or more user-provided input genes across the DepMap dataset. This is illustrated in Figure 2 using BRAF-mutated melanoma as an example. Entering BRAF as the input gene and applying filters for skin cancer, melanoma subtype, and BRAF-mutated cell lines restricts the analysis to a defined subset of 50 cell lines. The resulting network (**Fig. 2A**) reveals multiple genes that correlate with BRAF in this specific context. In this filtered analysis, *MAPK1* (encoding ERK2) shows the strongest correlation, followed by *MAP2K2* (encoding MEK2). This is consistent with the known biology of the MAPK pathway, in which ERK2 functions as a key downstream effector of BRAF signaling (23). It is also consistent with clinical practice, where MEK inhibitors (targeting the MEK1/MEK2 kinases encoded by *MAP2K1*/*MAP2K2*) are combined with BRAF inhibitors as standard of care in *BRAF*-mutated melanoma to suppress resistance mediated by MAPK pathway reactivation (24,25).

**Figure 2.**
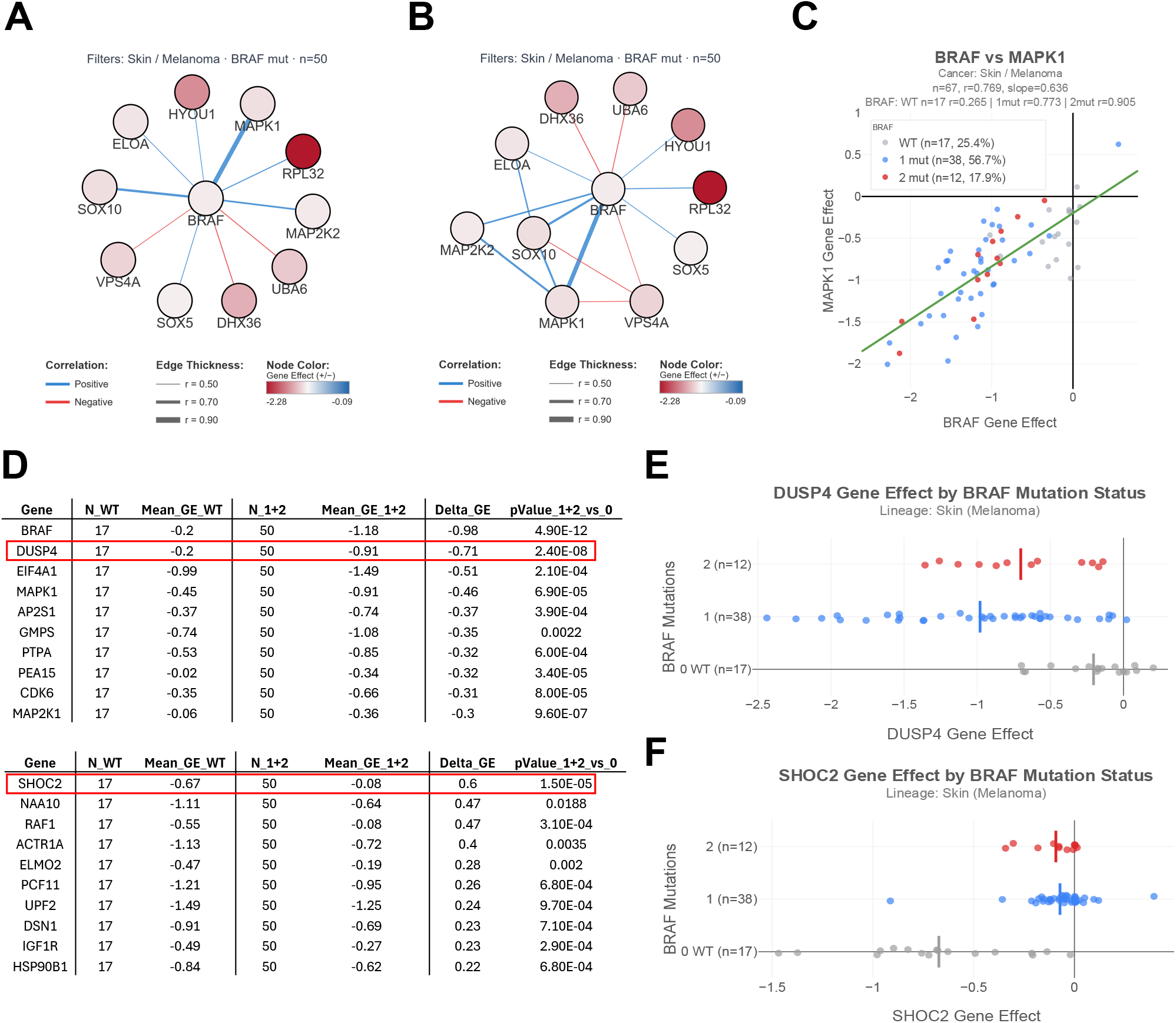
Hypothesis generation in Correlate. (**A**) Design mode network for *BRAF* with skin cancer/melanoma/*BRAF*-mutated filters (n=50 cell lines), with node colors reflecting gene effect scores (“Color by GE” mode). (**B**) Same network with the “Show all correlations” option enabled, revealing functional relationships among the correlated genes. (**C**) *BRAF* vs *MAPK1* scatter plot with *BRAF* hotspot mutation stratification. (**D**) Mutation analysis table showing differential gene dependencies between *BRAF* wild-type and *BRAF*-mutated skin cancer cell lines. (**E**) *DUSP4* gene effect stratified by 0, 1, or 2 BRAF hotspot mutations in melanoma. (**F**) *SHOC2* gene effect stratified by 0, 1, or 2 *BRAF* hotspot mutations in melanoma. A-C and E-F are directly exported from the app as SVG files. D is based on an exported .csv file.

Using the “Expand network” option in Design mode further allows the exploration of correlations among the identified genes (**Fig. 2B**). This is particularly useful for assessing whether genes identified by the initial query also correlate with one another, thereby revealing a broader co-dependency structure among them. Strong correlations among MAP2K2, MAPK1, and SO×10 are consistent with the published literature linking SOX10 suppression to tolerance to MAPK-targeting therapy (26,27).

**Figure 2C** shows the scatter plot for the *BRAF*-*MAPK1* correlation, the strongest in this analysis (r=0.77). The scatter plot can also be stratified by mutation status, here shown for *BRAF* hotspot mutations, with different colors indicating wild-type, single, or double mutation cell lines.

The analysis can be extended using the Mutation Analysis mode (selected in “1. Set Parameters”, indicated in Fig. 1A), which allows exploration of how specific hotspot mutations or translocations/fusions affect the gene effect of other genes across cell lines. Selecting *BRAF* as the hotspot gene with a skin cancer/melanoma filter generates a table showing genes whose gene effect differs significantly between BRAF wild-type and *BRAF*-mutated cell lines (**Fig. 2D**). This type of analysis can highlight potential drug targets in cancers carrying specific mutations. For example, *DUSP4* emerges as a gene with increasingly negative gene effect scores, indicating greater dependency, in melanoma cell lines carrying *BRAF* hotspot mutations (**Fig. 2E**). This is consistent with the known biology of DUSP4, a dual-specificity phosphatase that acts as a negative regulator of MAPK signaling and becomes essential when the pathway is constitutively activated by *BRAF* mutations (28,29). This mode can also reveal reduced dependencies in cancers carrying specific mutations. For example, *BRAF*-mutant melanoma shows significantly reduced, if not absent, dependency on SHOC2, an upstream regulator of the MAPK pathway (**Fig 2F)**.

This is consistent with the redundancy of upstream signaling imposed by the constitutively active *BRAF* V600E mutant (30). An equivalent analysis mode is also available for translocation/fusion events.

### Additional features

Correlate includes a synonym and ortholog lookup tool, available both as an integrated mode and as a standalone utility, that converts alternative gene names or mouse gene symbols to human gene symbols. This is particularly useful when analyzing data from mouse CRISPR screens, as the DepMap dataset is based on human cell lines. The application also provides embedded filters at multiple levels, such as “Compare by tissue” and “Compare by Hotspot/Fusion”, enabling context-specific analysis across cancer types and mutational backgrounds. A cell line browser allows users to search for cell lines by name and view associated metadata, including interactive filters for cancer type, subtype, and mutation status, thereby supporting the selection of appropriate models for experimental validation.

Correlate further supports downstream interpretation by allowing filtered gene lists to be directly queried for gene set enrichment against pathway resources such as KEGG and Reactome via the Enrichr API (31).

All network graphs, scatter plots, and bar charts can be exported directly from the application as SVG files suitable for use in vector graphics software such as Adobe Illustrator, Affinity Designer, or Inkscape. Correlate is designed as a companion to Green Listed v2.0, and gene lists explored in Correlate can be directly transferred to design custom CRISPR screen libraries (17).

## DISCUSSION

Correlate complements the existing landscape of DepMap analysis tools by providing an interactive, web-based platform for bulk correlation analysis of user-defined gene sets. While the DepMap portal offers powerful tools for exploring individual gene dependencies (4), and other tools such as shinyDepMap (14) and DepLink (15) provide gene essentiality browsing and drug-gene linkage, respectively, Correlate adds the ability to analyze gene lists in bulk, generate correlation networks, and explore hypothesis-driven questions in defined biological contexts, all directly in the browser without installation. The application builds on the R script for DepMap correlation analysis presented in Henkel et al. (17) and extends it with an interactive graphical interface and additional analysis modes.

Notably, Correlate’s approach is fundamentally different from knowledge-based methods such as GSEA (12) and STRING (13), which rely on annotated gene sets or known protein interactions. By instead identifying gene connections based on functional CRISPR screen readouts, Correlate can reveal relationships not present in curated databases, making it a complementary approach alongside established gene set and interaction analysis methods.

There are several constraints to consider when interpreting results. Specific mutations are often enriched in certain cancer types, and filtering by mutation or translocation/fusion status without also applying cancer type filters may result in comparisons that reflect lineage differences rather than the effect of the genetic alteration itself. For example, filtering for BRAF hotspot mutations without a cancer type filter would largely compare skin cancer cell lines against a mixed set of other lineages. Mutation and translocation/fusion filters should therefore be combined with cancer type and subtype filters to reduce such confounding. Additionally, the gene effect data in DepMap are based on genome-wide CRISPR knockout screens with survival and proliferation readouts, and the relationships identified by Correlate therefore reflect shared effects on cell fitness rather than direct physical or regulatory interactions. While this is highly relevant for cancer research, interactions that do not measurably affect cell survival or proliferation may not be captured with the same consistency.

Beyond correlation analysis, Correlate offers features that support both data visualization and experimental design. The ability to overlay user-provided CRISPR screen statistics on correlation networks (“With Stats” mode, “color by stats”) provides an integrated view of screen results and functional gene relationships. The essentiality overview (“color by gene effect”) can further support the design of custom CRISPR screen libraries. For example, broadly essential genes (very negative gene effect scores) could be excluded to create a more focused library, while the Design mode can be used to expand a library with correlating genes, adding granularity to a screen targeting a specific pathway. Importantly, the stats and essentiality overview is not only presented as a network but can also be generated and exported as a table, via the “Clusters” tab above the network. Correlate also facilitates cell line selection for experimental follow-up, as individual cell lines can be identified in scatter plots and explored through the cell line browser, including an interactive mutation filter via the “Oncoprint” button, helping users find appropriate cell models for their research questions. For example, users can rapidly identify TP53 wild-type cell lines harboring BCR translocations.

The application was designed to require no computational expertise, installation, or login, making it accessible to experimental biologists. It runs entirely client-side, ensuring responsiveness and avoiding reliance on server-side computation. However, because all computations are performed in the user’s browser, analyzing large gene lists in “Design mode” may be limited by the performance of the user’s computer. As the DepMap project continues to expand and release updated datasets, Correlate will be updated accordingly. Video tutorials demonstrating the use of Correlate and related tools are available through the Wermeling Lab YouTube channel.

## Acknowledgments

We acknowledge the Broad Institute Cancer Dependency Map project for making gene effect, mutation, and translocation/fusion data publicly available. We thank the Ma’ayan Laboratory for providing the Enrichr API. We are also grateful to Christer Uvehag, Daniel Uvehag, and CMM IT for assistance with server deployment and hosting infrastructure, and Dr. Martin Henkel and Prof. David P. Lane for suggestions.

## Author Contributions

S.D.: Investigation, validation, writing (review and editing). F.W.: Conceptualization, methodology, software, writing (original draft), visualization, supervision.

